## Supplemental Figure Legends for "Focal adhesion-derived liquid-liquid phase separations regulate mRNA translation"

### Supplementary Figure Legends

**Supplementary Figure S1.** (A) Comparison of disorder probability of *Homo sapiens* (human) and *Mus musculus* (mouse) p130Cas. Plot of disorder probability prediction versus amino-acid residue position in human (black) and mouse (red) p130Cas/BCAR1. Top: Color-coded protein domains. Horizontal line at 0.5 represents 5% false positive rate prediction. (B) Immunoblot of p130Cas in EGFP-p130Cas NIH3T3 cell line under transient over-expression. (C) Immunofluorescence image of transiently transfected NIH3T3 with EGFP-p130Cas (green) stained with anti-p130Cas antibody (red) showing colocalization. Arrows points to droplets. Scale bar = 20µm. (D-F) Inverted intensity images of EGFP-p130Cas transfected in (D) Chinese Hamster Ovary (CHO) cells (Scale bar = 50µm), (E) HeLa cells (Scale bar = 20µm) and (F) HEK 293Tx cells (Scale bar = 20µm). (G) Immunoblot of p130Cas in NIH3T3 and MCF7 cell line with GAPDH as a loading control. MCF7 cells express 2-2.5x more p130Cas than NIH3T3 fibroblasts. (H) Inverted intensity immunofluorescence image of endogenous p130Cas in MCF7 cells stained with anti-p130Cas antibody. Scale bar = 20µm. (I) Normalized histogram of p130Cas droplet area at ~6 hr (black) and ~24 hr (red) after plating transfected NIH3T3 fibroblasts on fibronectin-coated glass bottom dishes. N = 521 droplets from 48 cells and N = 631 droplets from 121 cells for 6 hr and 24 hr respectively.

**Supplementary Figure S2.** (A) Cell co-transfected with EGFP-p130Cas (green- left panel) and tagRFP-paxillin (red- middle panel) showing colocalization (merge- right panel). (B-C) Corresponding line intensity profiles for a typical droplet (B) and a FA (C). (D-E) Cell co-transfected with EGFP-p130Cas (green- left panel) and tagRFP-paxillin (D) or mCherry-FAK (E) (red- middle panel) showing colocalization in droplets with (lower panel) or without (upper-panel-control)) 250mM ammonium acetate treatment.

**Supplementary Figure S3.** Venn diagram showing the number of proteins significantly enriched in p130Cas droplets relative to GFP-only vs its published direct interactors.

**Supplementary Figure S4.** (A) HEK293T cells stably expressing EGFP-Dcp1A (left-panel) transfected with tagRFP-p130Cas (red-middle panel) and merged (right-panel). (B) A typical line intensity profile across a p130Cas droplet and a p granule.

**Supplementary Figure S5.** Immunofluorescence of Ago2 (A & B) and GW182 (C & D) (purple- middle panel) on GFP-Trap magnetic beads incubated with cell lysate from HEK293tx cells transfected with either EGFP-p130Cas (A & C) or EGFP (B & D) after fixation with 4% PFA. Corresponding DIC images, right panel.

**Supplementary Figure S6.** (A) NIH 3T3 cells on low or high FN labeled with puromycin (purple), nucleus labeled with hoeschst 33343. Cells were treated with or without 100µg/ml cycloheximide for 2 min before puromycin labelling for 10 min. (B) Quantification of p130Cas Western blot after knock-down in MCF7 cells. (C) HUVECs on low vs high FN labeled with puromycin as in A. (D) Quantification of puromycin labelling intensity in C. N = 388 and 348 cells from 9 field of views each from 3 independent experiments for low and high FN conditions respectively. (E) NIH 3T3 cells transfected with control or p130Cas siRNA on low vs. high FN, labeled for puromycin as in (A). (F) Immunoblot of p130Cas for NIH 3T3 cells in (E), GAPDH as loading controls. Labelling intensity quantified in (G). N = 4 field of views each from 4 independent experiments for each condition.

**Supplementary Figure S7.** (A-B) MCF7 cells expressing light-inducible cry2-tagRFP-p130Cas after exposure to 470nm LED light and stained for paxillin (A) or FAK (B) (green-middle panel) in (red- left panel), merge-right panel. Scale bar = 50µm. (C) MCF7 cells stably expressing cry2-tagRFP-p130Cas were sorted for modest vs high expressors and cells analyzed for tagRFP intensity. Top panel: untransfected MCF7 cells (negative control), unsorted cells (middle panel), sorted cells (modest over-expression -light blue) (bottom panel). (D) Immunoblot of p130Cas in WT and stable cry2-tagRFP-p130Cas expressing MCF7 cell line with GAPDH as a loading control.  (E) Pulse sequence with 5 sec ON followed by 60 sec OFF to minimize phototoxicity. Blue LED light array was controlled using programmable switch-Arduinos. (F). Puromycin labelled control WT MCF7 cells after 0- and 120-min exposure to blue LED light pulse protocol and (G) quantification of normalized puromycin intensity under these two conditions showing no effect of illumination on protein synthesis. N = 680 and 591 cells respectively for 0 and 120 min from 16 field of views each from 3 independent experiments.
