## Supplementary figures and images for "Focal adhesion-derived liquid-liquid phase separations regulate mRNA translation"

### Supplemental Fig S1

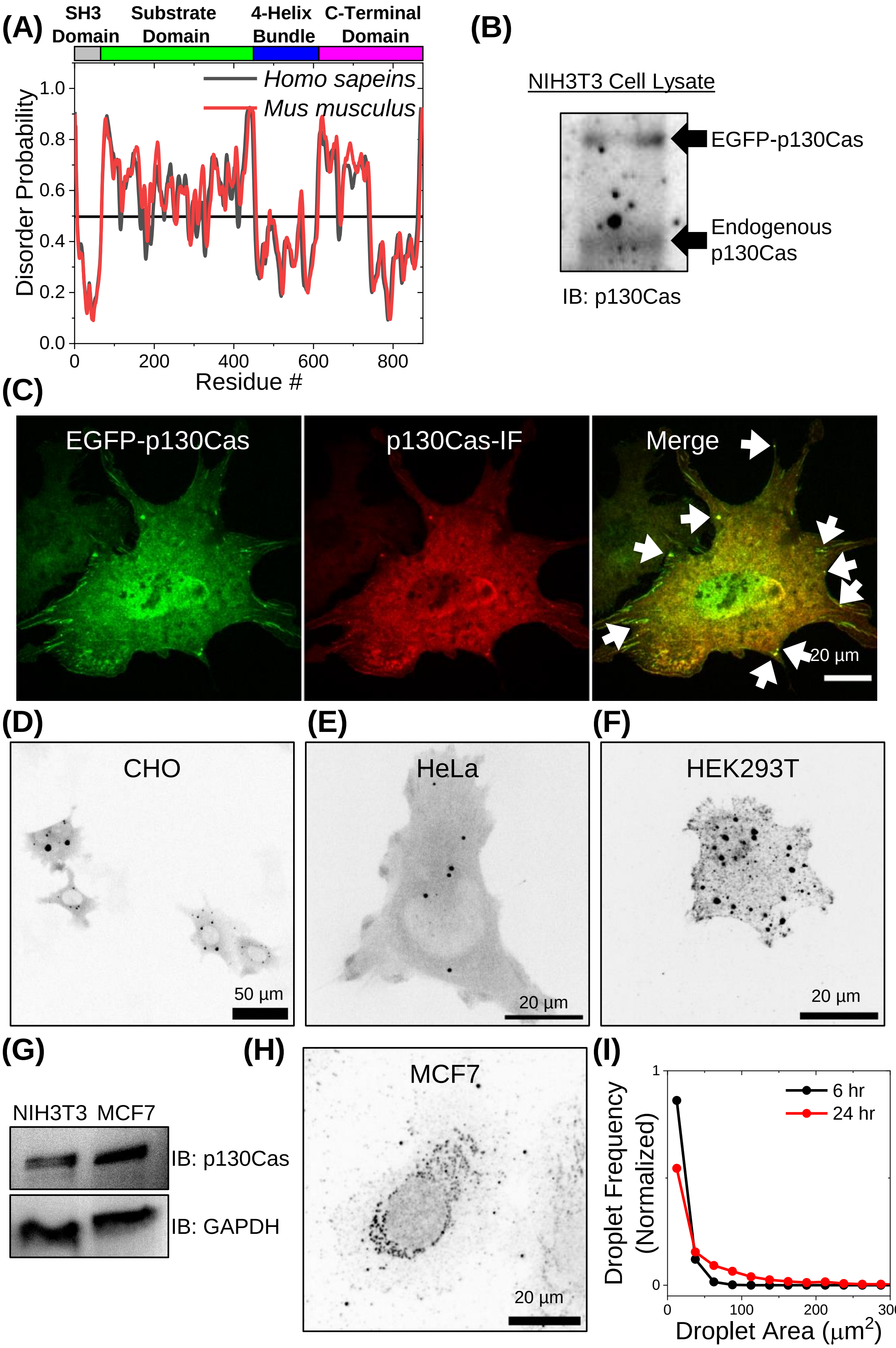

### Supplemental Fig S2

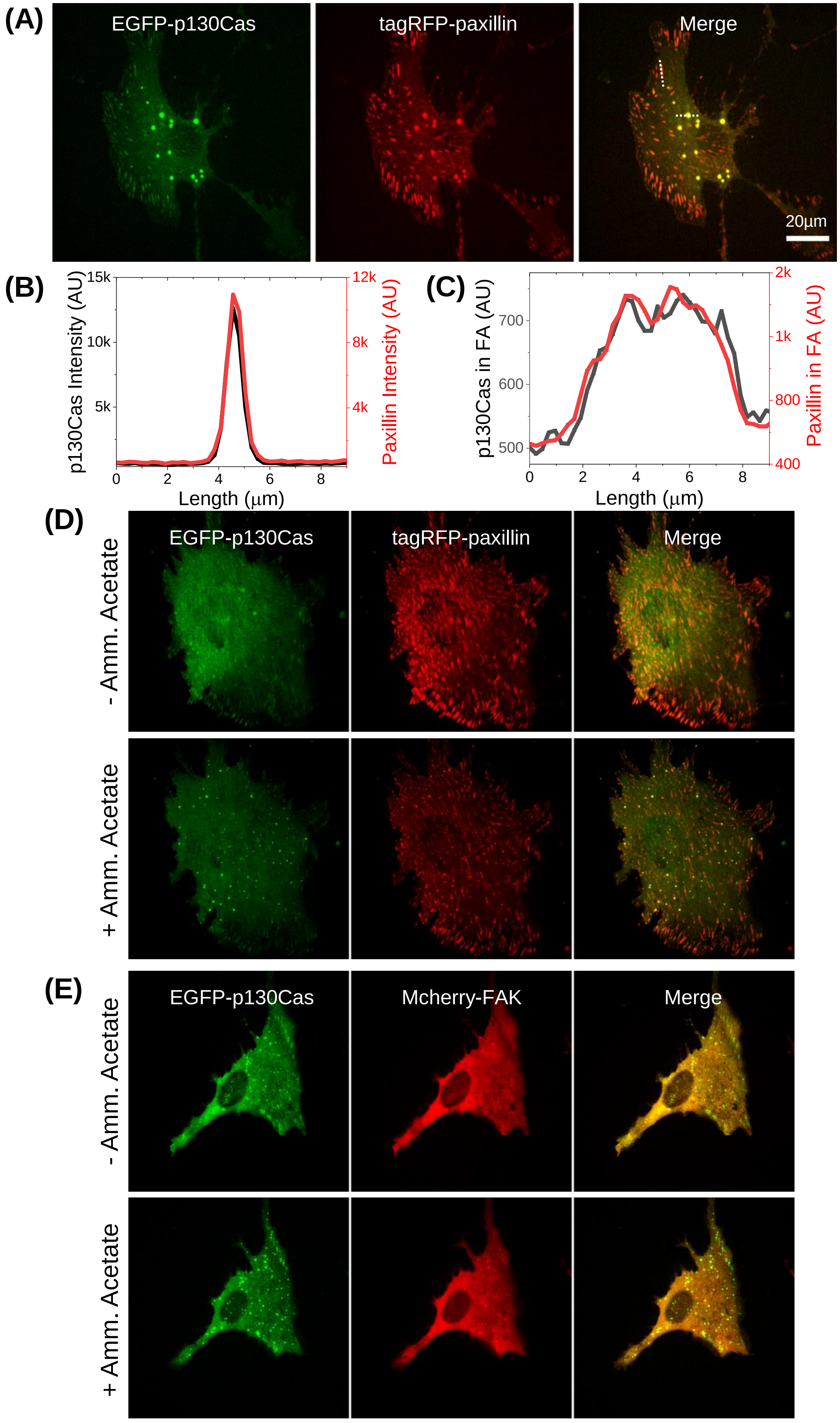

### Supplemental Fig S3

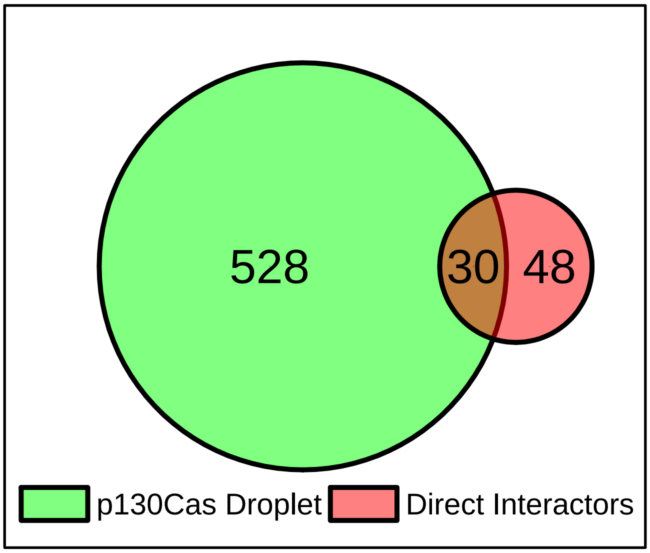


Figure S3

### Supplemental Fig S4

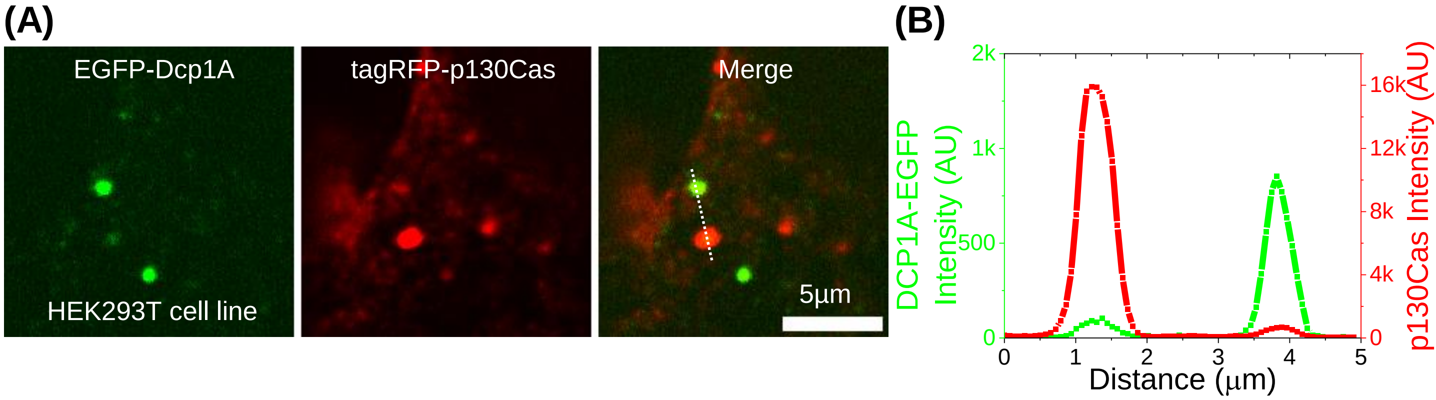

### Supplemental Fig S5

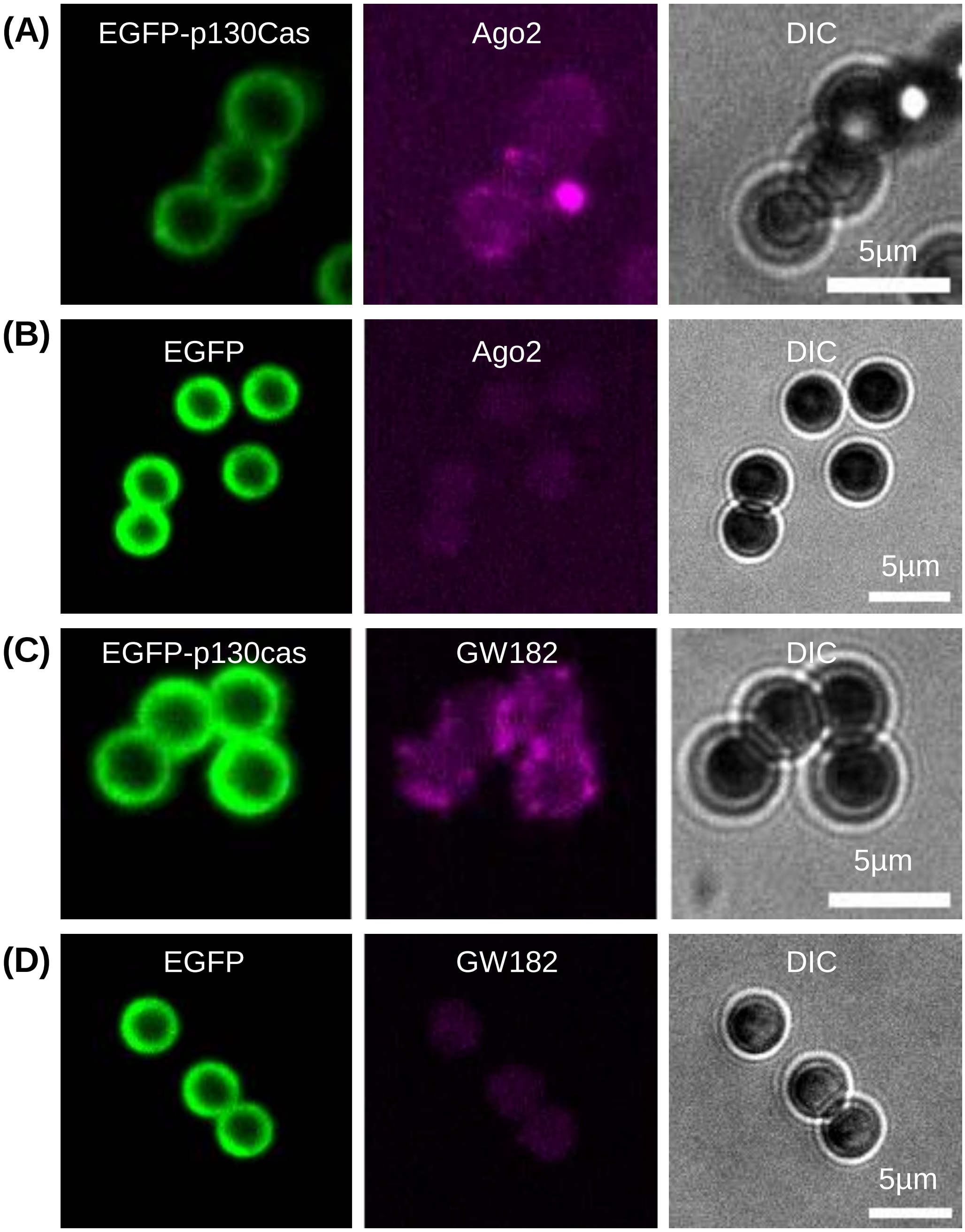

### Supplemental Fig S6

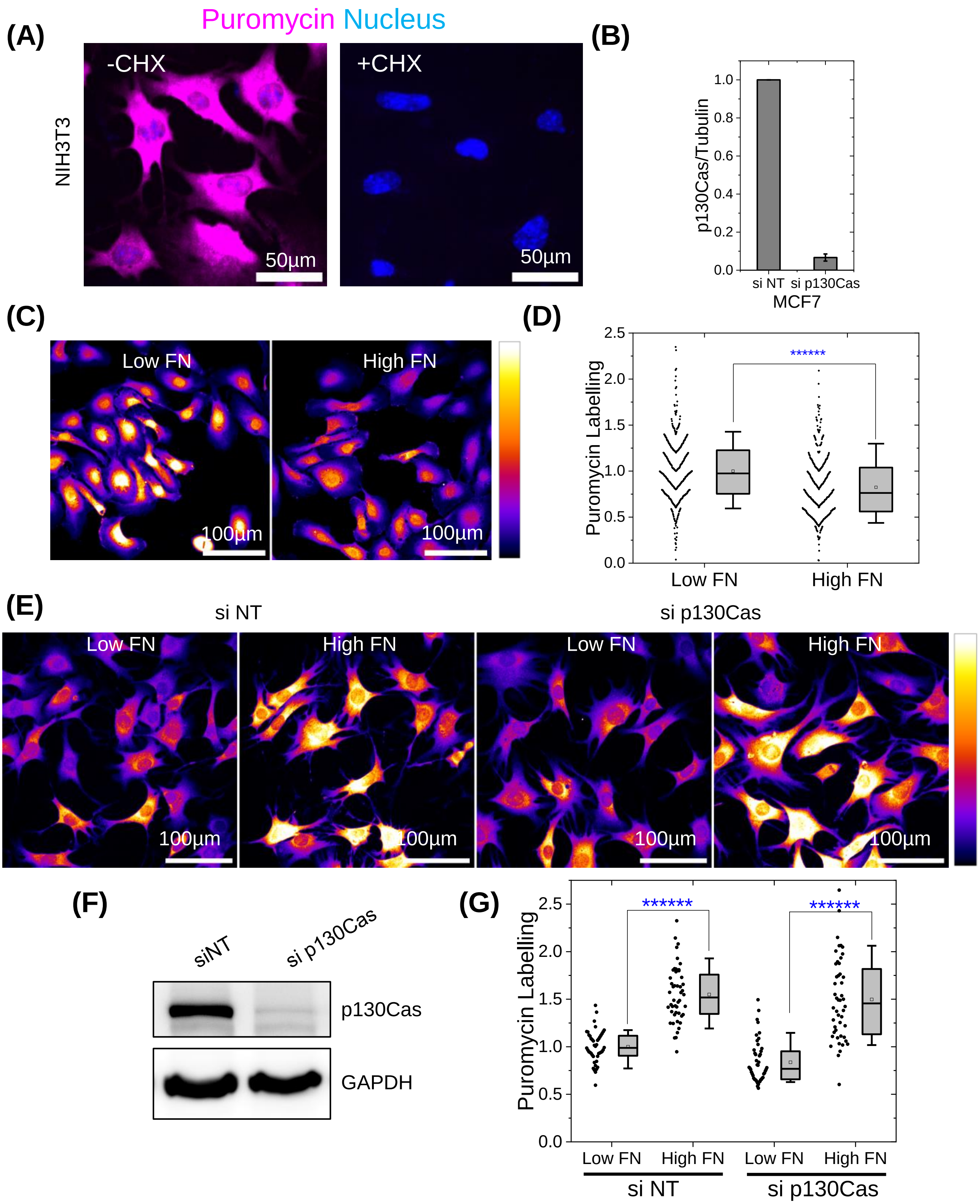

### Supplemental Fig S7

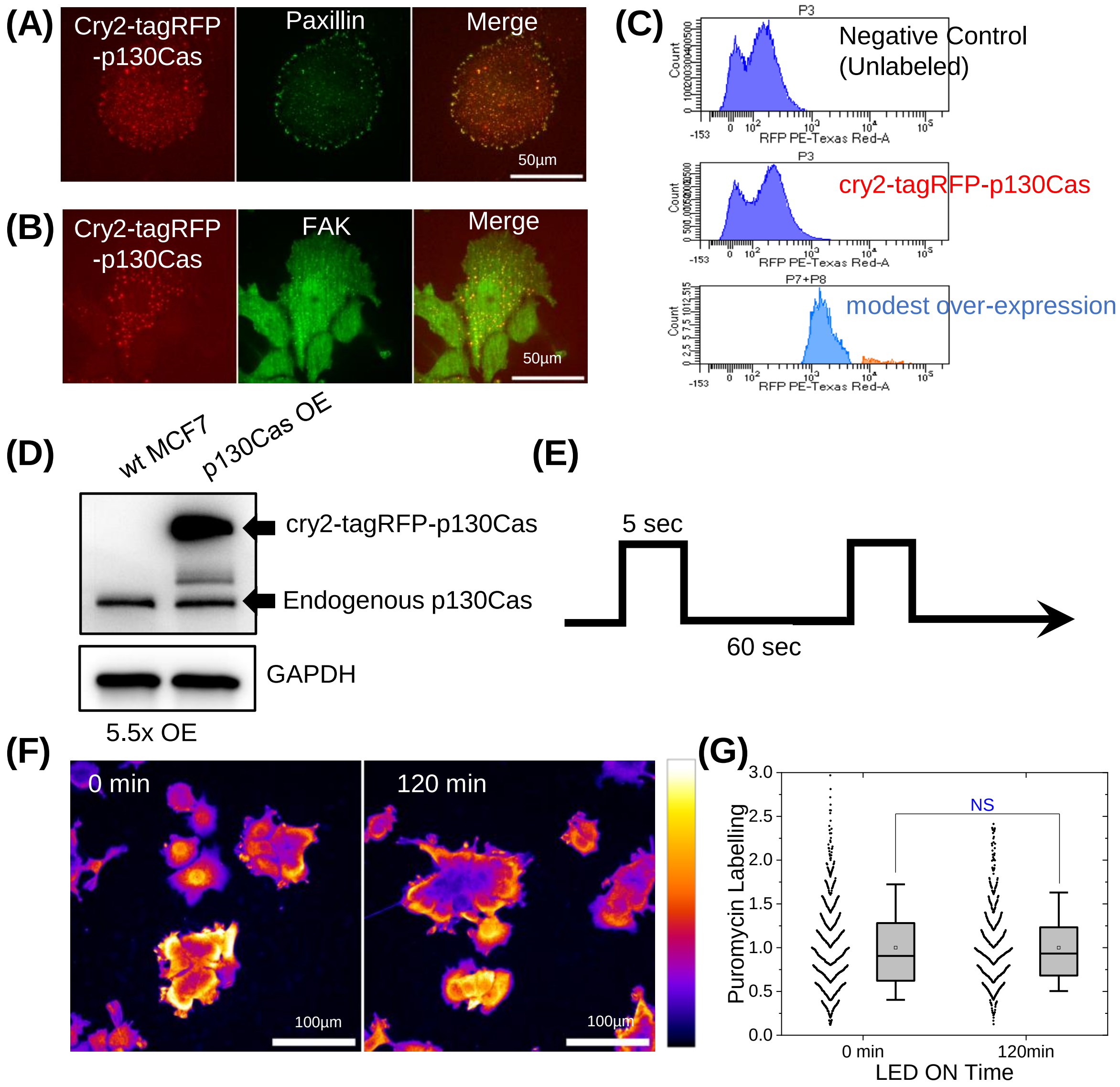
