## Supplemental Movie Legends for "Focal adhesion-derived liquid-liquid phase separations regulate mRNA translation"

### Supplementary Movie Legends

**Supplementary Movie S1.** Movie showing emergence of p130Cas droplets from FA in a live NIH3T3 fibroblast transiently transfected with EGFP-p130Cas freshly plated on fibronectin coated glass bottom dishes. Time points are shown in each frame, frame interval = 30 seconds. Scale bar = 10µm.

**Supplementary Movie S2.** Movie showing coalescence of two cytoplasmic p130Cas droplets in a live NIH3T3 fibroblast transiently transfected with EGFP-p130Cas freshly plated on fibronectin coated glass bottom dishes. Time points are shown in each frame, frame interval = 30 seconds. Scale bar = 10µm.

**Supplementary Movie S3.** Movie of p130Cas droplet merging back into a FA in a live NIH3T3 fibroblast transiently transfected with EGFP-p130Cas freshly plated on fibronectin coated glass bottom dishes. Time points are shown in each frame, frame interval = 30 seconds. Scale bar = 10µm.

**Supplementary Movie S4.** Movie of p130Cas droplet co-emerging from FA along with paxillin. EGFP-p130Cas is in green and tagRFP-Paxillin is in red. Time points are shown in each frame, frame interval = 2 minutes. Scale bar = 10µm.

**Supplementary Movie S5.** Movie of p130Cas droplet co-emerging from FA along with FAK. EGFP-p130Cas is in green and mcherry-FAK is in red. Time points are shown in each frame, frame interval = 1 minutes. Scale bar = 10µm.

**Supplementary Movie S6.** Movie of induction of p130Cas droplets in MCF7 cell line stably expressing modest levels of cry2-tagRFP-p130Cas upon irradiation with blue laser light for 0.5 second. 0 second frame is before irradiation and 15 second onwards frame is after irradiation. Time points are shown in each image, frame interval = 15 seconds. Scale bar = 50µm.
