## Supplemental Table legends for "Focal adhesion-derived liquid-liquid phase separations regulate mRNA translation"

### Supplementary Table Legends

**Supplementary Table 1.** Excel Sheet containing the list of proteins found in p130Cas droplets determined by mass spectrometry.

**Supplementary Table 2.** Excel Sheet containing the list of mRNAs found in p130Cas droplets determined by RNA sequencing.

**Supplementary Table 3.** Excel Sheet containing the list of differentially regulated mRNAs in HEK293T cells with p130Cas droplets induced by its overexpression as determined by RNA sequencing.
